# *matchRanges*: Generating null hypothesis genomic ranges via covariate-matched sampling

**DOI:** 10.1101/2022.08.05.502985

**Authors:** Eric S. Davis, Wancen Mu, Stuart Lee, Mikhail G. Dozmorov, Michael I. Love, Douglas H. Phanstiel

**Author notes:** **Availability and implementation** https://nullranges.github.io/nullranges. These authors co-supervised the work (Michael I. Love, Douglas H. Phanstiel). Contact: Douglas H. Phanstiel, Michael I. Love.

## Abstract

Deriving biological insights from genomic data commonly requires comparing attributes of selected genomic loci to a null set of loci. The selection of this null set is non trivial, as it requires careful consideration of potential covariates, a problem that is exacerbated by the non-uniform distribution of genomic features including genes, enhancers, and transcription factor binding sites. Propensity score-based covariate matching methods allow selection of null sets from a pool of possible items while controlling for multiple covariates; however, existing packages do not operate on genomic data classes and can be slow for large data sets making them difficult to integrate into genomic workflows. To address this, we developed *matchRanges*, a propensity score-based covariate matching method for the efficient and convenient generation of matched null ranges from a set of background ranges within the Bioconductor framework.

## Introduction

Genome-wide analyses can provide valuable insights into biological systems and human disease by revealing patterns of features that may be missed by interrogation of individual loci. Determining if observed trends are statistically significant, however, commonly requires comparing attributes between a focal and a null set of genomic loci. Accurate inference requires that null sets exhibit similar distributions of covariates observed in the focal set, to mitigate interpretability issues due to confounding. This can be challenging since many common covariates (e.g., GC content, gene density, histone acetylation, chromatin accessibility, etc.) are not uniformly distributed throughout the genome and must therefore be explicitly controlled when selecting null sets of loci^1^. Propensity score-matching is a computational method that allows for the selection of covariate-matched sets and several packages implement it within the R programming language^2,3^. However, these packages can be slow for genome-scale data sets and are not well-integrated into genomic analysis platforms such as Bioconductor making them difficult to incorporate into genomic workflows.

To address this problem, we developed *matchRanges*, an efficient and convenient tool for generating covariate-matched sets of genomic ranges from a pool of background ranges. *matchRanges* computes for each range a propensity score, the probability of assigning a range to focal or background groups, given a chosen set of covariates. It provides three methods including nearest-neighbor matching, rejection sampling, and stratified sampling for null set selection^4^. Additionally, *matchRanges* provides utilities for accessing matched data, assessing matching quality, and visualizing covariate distributions. The code has been optimized to accommodate genome scale data sets, such that most matchRanges functions can efficiently process sets of millions of loci in seconds on a single core (**Fig S1**). *matchRanges* accepts and returns common Bioconductor objects, such as *GRanges* and *GInteractions* for seamless integration with existing workflows^5–7^ (**Fig S2**). matchRanges is distributed as part of the *nullranges* package, with multiple software vignettes.

### The *matchRanges* workflow

To generate a covariate-matched set of ranges, users can provide *data*.*frame, GRanges* or *GInteractions* R objects annotated with columns describing one or more potentially confounding covariates^6–8^. The *matchRanges* function takes as input a “focal” set of data to be matched and a “pool” set of background ranges to select from. *matchRanges* performs subset selection based on the provided covariates and returns a null set of ranges with distributions of covariates that approximately match those of the focal set (**Fig 1A**). This allows for an unbiased comparison between features of interest in the focal and matched sets without confounding by matched covariates. As the returned matched sample object is the same class as the inputs, it can be easily incorporated into new or existing Bioconductor workflows^9^.

**Fig. 1.**
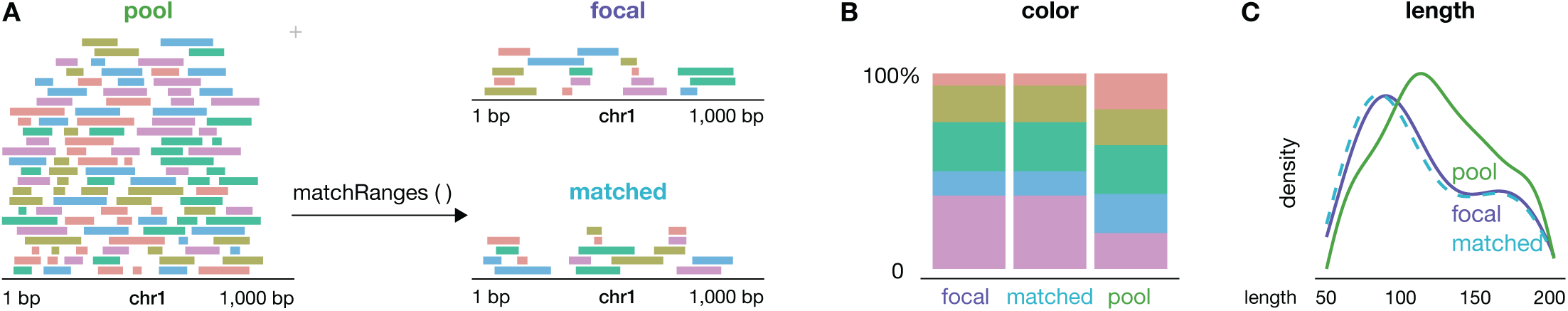
matchRanges workflow. **(A)** A schematic demonstrating how the *matchRanges* function can be used to select a set of *GRanges* matched for covariate features of color and length. **(B and C)** Example visualization of covariate distributions for assessing matching quality and covariate balance. Figure generated with the *plotgardener* R/Bioconductor package^11^.

A key aspect of inference based on covariate matching is visual inspection of the results. We provide several functions to assess the overall quality of matching, including plots of the distribution of covariates amongst the “focal”, “pool”, and “matched” sets (**Fig 1B and C**). Accessor functions allow users to easily extract data for further inspection or integration with covariate balance packages, such as *cobalt*^*10*^. Since matching is a pre-processing step, multiple matching methods can be tried and assessed before downstream analyses.

Detailed documentation on how to use *matchRanges* and when to use each matching method is available at an accompanying website (https://nullranges.github.io/nullranges), which contains step-by-step tutorials and biological case studies demonstrating the power of *matchRanges*.

## Conclusion

*matchRanges* is a collection of R functions for generating covariate matched ranges to test associations between sets of genomic ranges. Distributed as part of the *nullranges* R package, *matchRanges* uses a propensity score-based method to perform subset selection on genomic ranges, allowing fair comparisons between two sets of interest while avoiding problems with confounding by nuisance covariates. The package provides functions for assessing, visualizing, and extracting matched data that integrates seamlessly into existing Bioconductor workflows. *matchRanges* will be useful to genomic researchers from all disciplines and will help accelerate scientific progress by improving the accuracy and rigor of genomic analyses.

## Acknowledgments

We thank Tim Triche and Kasper Hanson for their helpful feedback and suggestions.

## Supplementary information

Companion data package: https://github.com/nullranges/nullrangesData

Companion website: https://nullranges.github.io/nullranges

Code for figures: https://github.com/EricSDavis/matchRangesManuscript

**Supplementary Fig. 1.**
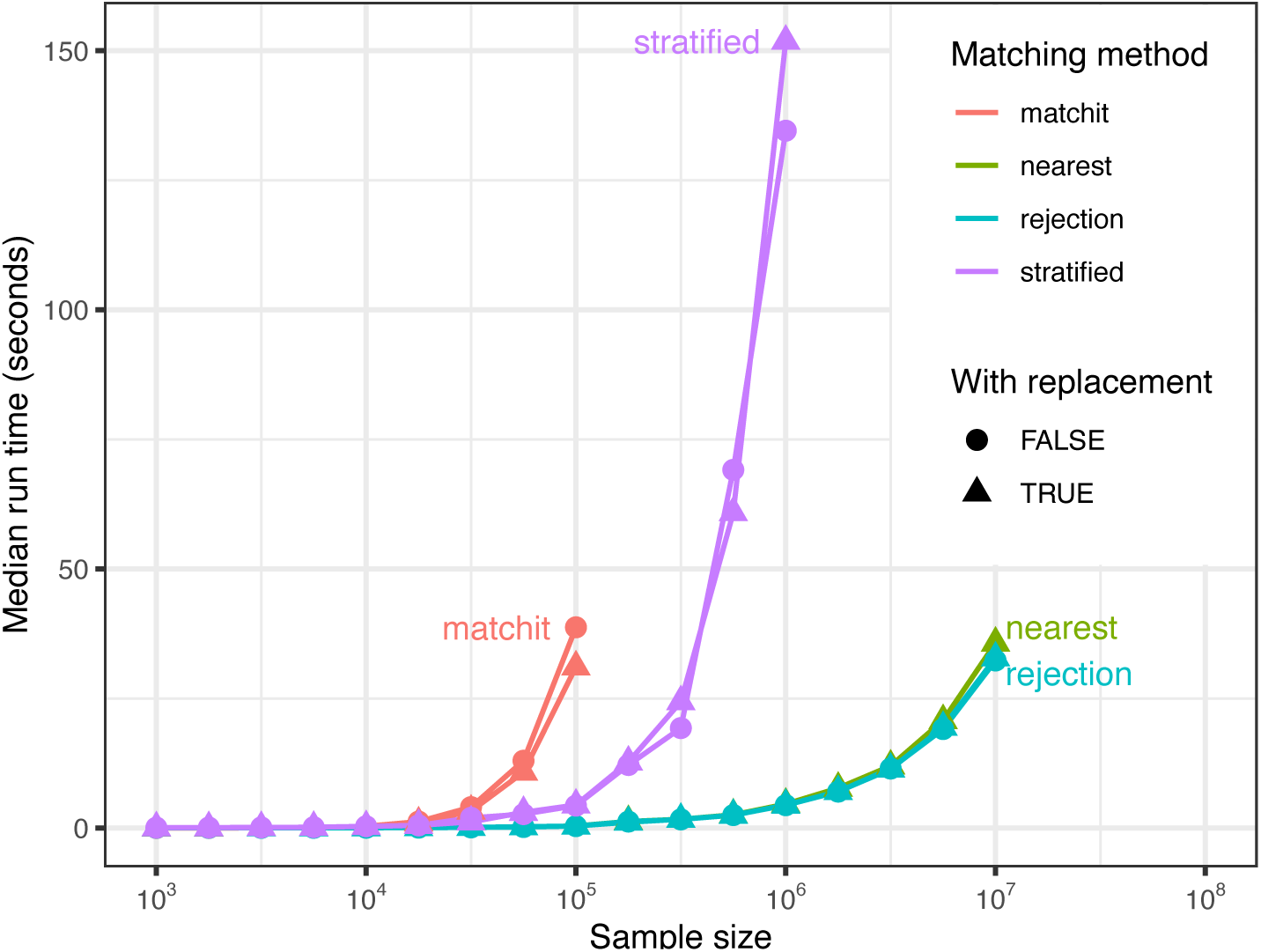
matchRanges run time. Median run time for *matchRanges and MatchIt* applied to simulated data. Data was matched for one continuous feature for each matching method and replacement option. Sample size contains 95% values as pool and 5% as focal. *MatchIt* was run with default parameters using method=“nearest”. Each point represents the median run time for n=20 points.

**Supplementary Fig. 2.**
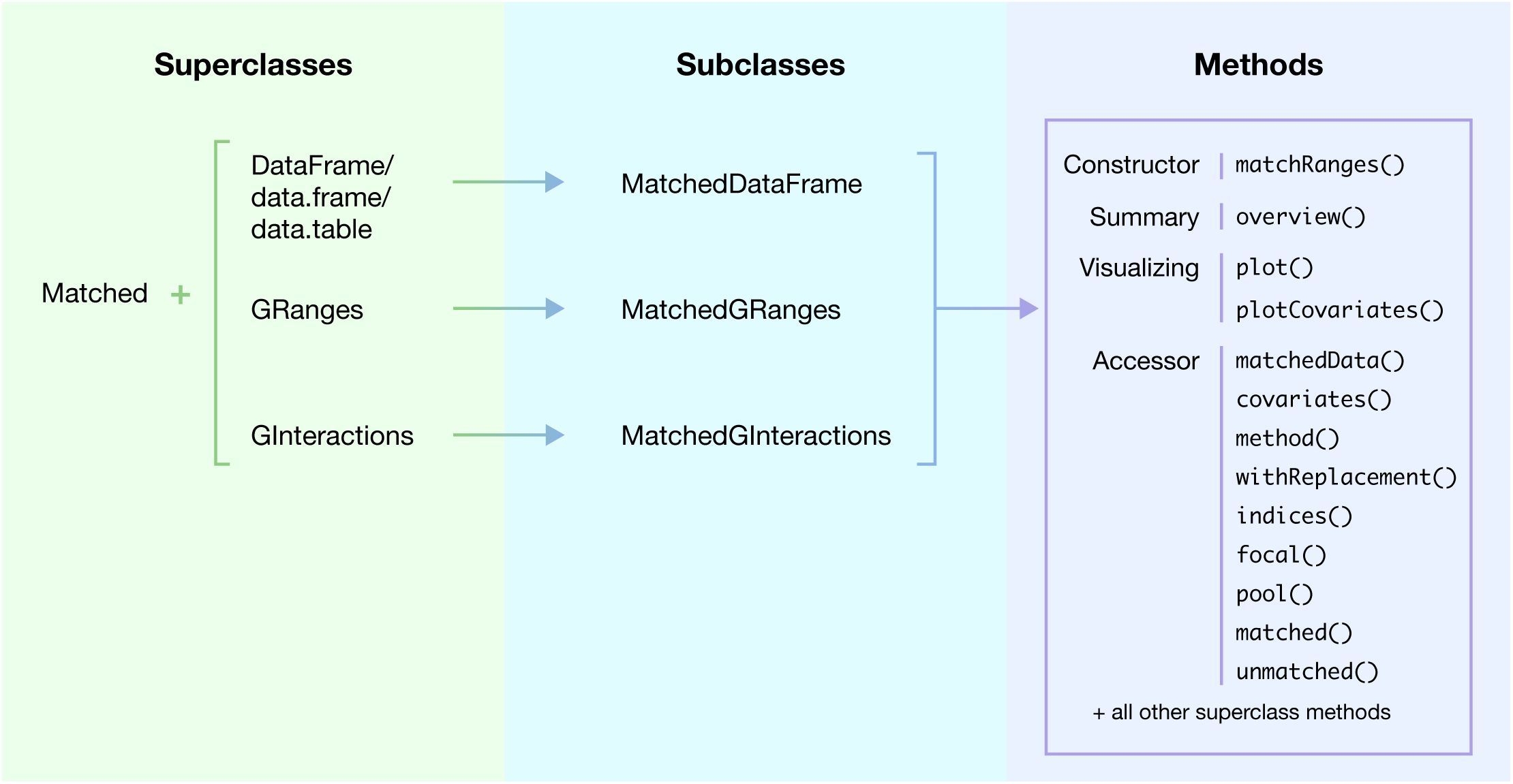
Class structure. Overview of the *matchRanges* class structure and methods. The *Matched* class is combined with either the *DataFrame, data*.*frame, data*.*table, GRanges*, or *GInteractions* classes (*left panel*) to create the *MatchedDataFrame, MatchedGRanges*, or *MatchedGInteractions* subclasses (*middle panel*). Each subclass behaves as a combination of both its superclasses - with access to both methods of the *Matched* class (*right panel*) and each respective class’ methods.

